# Local Generation and Efficient Evaluation of Numerous Drug Combinations in a Single Sample

**DOI:** 10.1101/2022.12.13.520254

**Authors:** Vlad Elgart, Joseph Loscalzo

## Abstract

We develop a method that allows one to test a large number of drug combinations in a single cell culture sample. We rely on randomness of drug uptake in individual cells as a tool to create and encode drug treatment regimens. A single sample containing thousands of cells is treated with a combination of fluorescently barcoded drugs. We create independent transient drug gradients across the cell culture sample to produce heterogeneous *local* drug combinations. After incubation period, the ensuing phenotype and corresponding drug barcodes for each cell are recorded. We use these data for statistical prediction of the response to the drugs treatment in a macroscopic population of cells. To further application of this technology, we developed a fluorescent barcoding method that does not require any chemical drug(s) modifications. We also developed segmentation-free image analysis capable of handling large optical fields containing thousands of cells in the sample, even in confluent growth condition. The technology necessary to execute our method is readily available in most biological laboratories, does not require robotic or microfluidic devices, and dramatically reduces resource needs and resulting costs of the traditional high-throughput studies.

## Introduction

The optimization of targeted therapies is based on the determination of drug dose-response and the selection of a regimen that is best suited for an individual patient. A critical barrier to optimization screening is the availability and amount of relevant tissue sample (e.g., biopsy), especially in the case of combination therapy. This limitation is a reflection of the fact that the number of possible drug combinations grows exponentially with the number of drugs and dosages (cf., Supporting Information). For example, the number of all possible combinations of three drugs tested at ten different concentrations (including zero) is 10^3^. This number does not include the biological replicates and controls that render the careful testing of even three drugs extremely challenging.

There exists another challenging aspect to the identification of optimal drug dosage and regimen: accurate reproducibility of *in-vivo* and *in-vitro* experiments is notoriously difficult to achieve in many biological applications, such as drug dose-response quantification. Many confounding factors contribute to the variability in sample readout [1, 2]; these factors can generally be grouped into two categories, which are usually described as intrinsic and extrinsic noise [3, 4]. Sources of intrinsic noise in drug uptake by and action within cells include fundamental biochemical processes, such as passive uptake, active transport, and degradation. By contrast, external factors that contribute to noise are principally manifest through convection.

While intrinsic cell variability can be significant, we believe that it is the extrinsic factor(s) that drive sample variability in most experimental cellular systems. This conclusion derives from the Law of Large Numbers [5], since the typical number of cells in a culture sample is at least of the order of 10^4^, while the number of biological/technical replicates is ~ 3.

If the observed sample variability is due to extrinsic noise, then it is very tempting to conduct all drug response experiments in the same tissue or cell culture sample. Indeed, under these conditions, it is guaranteed that external fluctuations in the environment are exactly the same for all testing conditions. Hence, quantification of the drug’s response does not require averaging over many different samples to achieve an accurate, *relative* comparison of the drug’s effectiveness.

In order to test different drug combinations in the same cell culture or tissue specimen, one must create these combinations *locally*. One of the smallest possible quantifiable locales in both cell culture and tissue samples is a single cell. Indeed, we can treat each cell as an individual chemical “reactor” in which testing of different drug combinations is performed. Thus, if we can (i) create different drug combinations in individual cells, and (ii) subsequently quantify these combinations and the ensuing outcome (phenotype) for each cell, the logistics of testing combinations in a single macroscopic sample would be effectively addressed.

There exists a straightforward way to resolve the quantification issue (ii) above. Drugs can be uniquely labeled (e.g., using fluorescent tags), in which case, the amount of drug taken up by each cell would scale as the fluorescence of its corresponding tag.

Let us now address a more delicate problem (i): how to create a broad range of drug combinations in different individual cells. At first glance, an herculean effort would be required to accomplish this task. Fortunately, cells themselves solve this problem effort-lessly for us. Even cells with identical DNA typically exhibit deviations in drug uptake. Therefore, the random nature of drug(s) uptake by individual cells can be utilized to create different drug combinations. Here, we further enhance heterogeneity and independence in *stochastic* drugs mixing by creating individual drug gradients across the sample.

Rather than delivering drugs homogeneously, we rely on simple forced drug convection (by local injection) to create transient local drug gradients independently for each drug. After delivery and incubation of random *local* drug combinations to individual cells, we analyze the ensuing cellular phenotypes and corresponding drug optical barcodes by fluorescence microscopy or flow cytometry. The statistical analysis of the imaging data is used to predict the effect of ratiometric dosing on the outcome (cell phenotype). In this fashion, we require as little as a single cell culture sample to test all feasible drug combinations, bypassing the need for an otherwise enormous number of controls and technical replicates.

In this work, fluorescently labeled drugs are clustered into three “barcoding” classes distinguished by the physical association of the label with the drug: either irreversibly linked or cleavable/separable chemistry. Additionally, we developed a labeling method that does not require any chemical modification of the drug(s), such as covalent fluorescent tagging. This particular approach involves local co-injection of an unconjugated pair of pre-mixed drug and fluorescent dye that serves as a correlative barcode. We utilize the fact that on short incubation time scales (less than an hour), the initial injection producing forced convection is mainly responsible for transient gradients in the sample. Immediately after local injection, the drug and its tag concentration profiles across the sample correlate. The subsequent diversion of these profiles due to differences in diffusion and transport rates between the drug and its unconjugated tag occurs slowly [6] over this comparatively short incubation period, ensuring persisting correlation between drug and tag uptakes throughout the course of the typical experiment. For all drug barcoding classes we designed and tested, we rely on the fact that the drug barcode is retained by the cells. Hence, total drug exposure over time (area under the curve) can be assessed by a single time point measurement.

We also developed segmentation-free microscopy-based image analysis and statistical data processing capable of handling large optical fields containing thousands of cells in the sample, even in confluent growth condition. These algorithms allow us to bypass the standard segmentation process, typically requiring additional fluorescent marker(s), significant computational time, and resources.

Lastly, we include real-world application(s) of our stochastic drug mixing strategy. Specifically, we conducted multiple cell culture experiments treating cells with combinations of drugs commonly used in multi-drug chemotherapy treatments, such as cisplatin, docetaxel, cyclophosphamide, and 5-fluorouracil. We adopted our technology to accommodate quantification of the cell death as a phenotype common to all of these agents, demonstrating the method’s applicability to practical challenges in combination drug development.

Our approach has distinct advantages over conventional drug combination screening methods, including far greater efficiency, a smooth (continuous) variation of drug concentrations in the resulting combinations, and concomitant phenotype readout.

## Main

### Stochastic Drug Mixing

We first consider a benchmark system in which “drugs” are diester dyes with distinguishable fluorescent properties. These dyes are well retained by the cell for an extended period and, hence, can be used to characterize total uptake by individual cells. Typical targets of these “drugs” are intracellular thiol or amine groups (Methods).

In Figure 1a we demonstrate the typical results of a random drug mixing experiment using a homogeneous delivery method. HeLa cells were exposed to four different dyes mixed together simultaneously in suspension cell culture according to the manufacturer’s protocol and imaged using confocal microscopy. The color of each dye (violet, green, yellow, and red) corresponds to its specific fluorescence emission maximum. The color and intensity of each cell in the image correspond to a unique “drug” combination.

**Figure 1:**
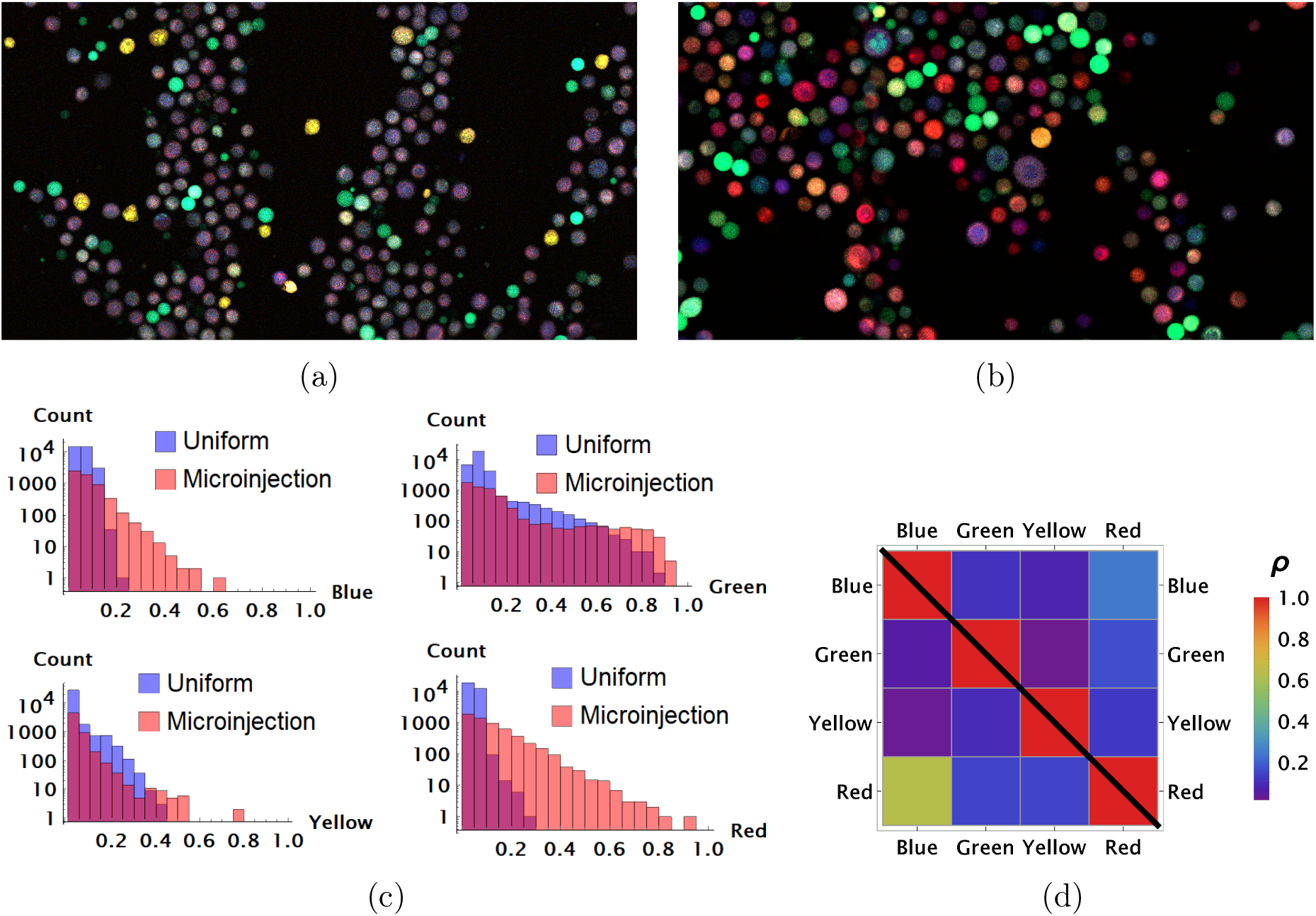
(a) and (b), Results of random dye mixing in suspension culture; raw image data are shown for homogeneous and point injection delivery methods, respectively. Here, we used four different CellTrace dyes (Violet, CFSE, Yellow, and Far Red), labeling sequentially as for dyes *A* and *B* in Figure 10. (c) The uptake distribution of individual CellTrace dyes incubated with HeLa cells in suspension is shown for each of the conditions on the same plot. Here, each histogram displays cell count as a function of fluorescence intensity, measured in arbitrary units for each dye. (d) Correlation coefficients, *ρ*, for all pairwise dye combinations are shown graphically for homogeneous and point injection deliveries on lower and upper triangular matrix plots, respectively. The absolute value of the correlation coefficient is represented by colors varying from violet to red for values 0 and 1, respectively.

One reason as to why the majority of cells exhibit somewhat similar color is the high degree of similarity in chemical and, hence, transport properties of these compounds. There is a propensity for cells to take up similar molecules proportionally, with only atypical cells (outliers) exhibiting different behavior in this case.

We experimented with multiple strategies for increasing variability, such as decreasing incubation time and cell culture volume, factors that we showed previously drove heterogeneity in drug uptake [7]. We found that the most robust way to control variability in uptake is to generate transient drug gradients in culture. A typical outcome of these experiments with HeLa cells is shown in Figure 1b as a raw image. Here, we used a rather crude and simple method to establish transient spatial gradients in suspension cell culture by point injection (Methods).

The quantitative comparison of homogeneous and point injection-based dye delivery in terms of the fluorescence intensity distribution and pairwise correlation relationship is shown in Figure 1c and Figure 1d, respectively. Additional heterogeneity introduced by point injections leads to a widening of the fluorescence intensity distributions and, thus, a significantly broader concentration range compared to the homogeneous mixing experiment.

We next conducted parallel experiments with adherent HeLa cells using gradient mixing method by point injection of different drugs in different locations in the culture well (cf. Methods).

Due to the dispersion process, the initial drug(s) distribution is spread (“averaged”) locally across finite regions of space (spread size depends on drug diffusion and uptake rates, and also incubation time). This observation allows one to simplify the image processing by averaging fluorescence signal intensities not over individual cells, but, rather, over the mesoscopic regions of space containing multiple cells (Methods). This simplification is critical since confluent cell growth conditions (necessary for a large statistical ensemble) usually present a problem for image segmentation. The statistics obtained using this approach describes typical local cell drug uptake within the given mesoscopic region (tile).

The results of a typical experiment utilizing point injections are compared to homogeneous dye delivery in Figure 2. Here, we either co-delivered all dyes homogeneously or locally by point injection. Point injections were performed by local delivery of two dye pairs simultaneously: One pair of *pre-mixed* dyes was codelivered in one point location and another pair of *pre-mixed* dyes was injected at a different point location on the microscopy cell culture slide. As in the case of cells grown in suspension culture, point injection greatly increased the range of drug uptake across the population compared to homogeneous delivery (Figure 2a).

**Figure 2:**
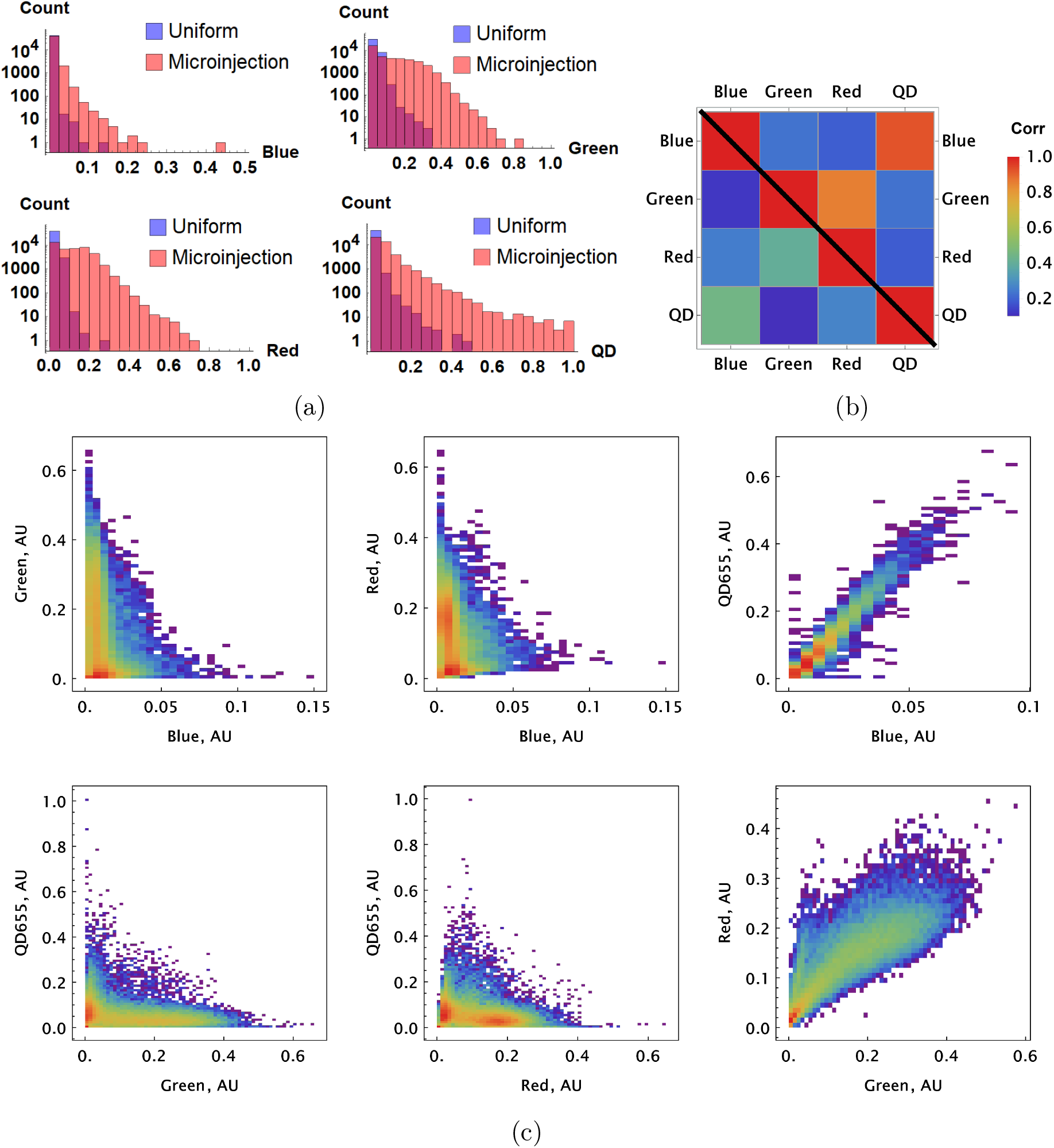
A pair of Green and Red CellTrace dyes was co-delivered in one point location and another pair of Blue and QD_655_ dyes was co-delivered in another point location. (a) and (b), Statistical properties of homogeneous and point drug injections (microinjection) are compared in the same fashion in Figures 1c and 1d. (c) Scatter density plots for drug uptakes are shown for each pair of dyes tested. First two columns, independent delivery of each dye pair; last column, co-delivery of each pair.

We observed correlation patterns between dye uptakes that were dependent on whether dyes were paired by point delivery or not. Locally co-delivered dyes (point-injection) exhibited a far greater degree of correlation compared to those in unmatched pairs of drugs delivered to different slide locations (Figure 2b). This behavior is apparent from the scatter plots shown in Figure 2c where color represents local cell count (higher count corresponds to a redder color in the color spectrum scheme). Dye pairs delivered independently by the point-injection method (two different culture slide locations) led to the generation of different mixing ranges of dyes (cf., first two columns, Figure 2c).

The pre-mixed dyes co-delivered locally exhibit a high degree of correlation with coefficients of correlation greater than 0.8 (cf., right column in Figure 2c), even for structurally and chemically different agents such as CellTrace and nanoparticle QD dyes. The underlying cause of this surprising behavior is identical *initial* dispersion of pre-mixed co-delivered dyes throughout a sample by forced convection. The subsequent diversion of concentration profiles due to differences in dye diffusion, transport, and reaction rates occurs slowly over this comparatively short incubation period, ensuring persisting correlation between dye uptakes throughout the course of the typical experiment.

### Conjugation-free Drug Barcoding

One can think of one of the dyes in co-delivered pairs as a drug and another as a label/tag. Since we can predict drug uptake utilizing the high degree of correlation with the corresponding tag, no physical conjugation of drug and tag is necessary.

Unlike chemically labeled drugs (where the one-to-one correspondence between tag fluorescence and drug uptake is assured by design), the conjugation-free barcoding method requires statistical processing. This step is necessary to filter out the noise due to intrinsic fluctuation in both drug and tag uptakes and improve correspondence between tag fluorescence and drug uptake.

A demonstration of the predictive accuracy in estimating drug uptake using this conjugation-free approach is shown in Figure 3. Here, coarse-grained uptakes of co-delivered dye pairs are shown in the scatter plots, Figure 3a and Figure 3b, with linear model fit and confidence bands depicted by blue and yellow lines, respectively. Average intensity values for the homogeneous dye delivery method are depicted by the large, red-filled circle on both graphs, demonstrating the greatly expanded range of combined dye concentrations by the point injection method. These results are also independent of the specifics of injection point location and geometry, as demonstrated in Figure 3c and Figure 3d, where we compare raw intensity data from two dyes in two different experiments involving two different pairs of injection site locations.

**Figure 3:**
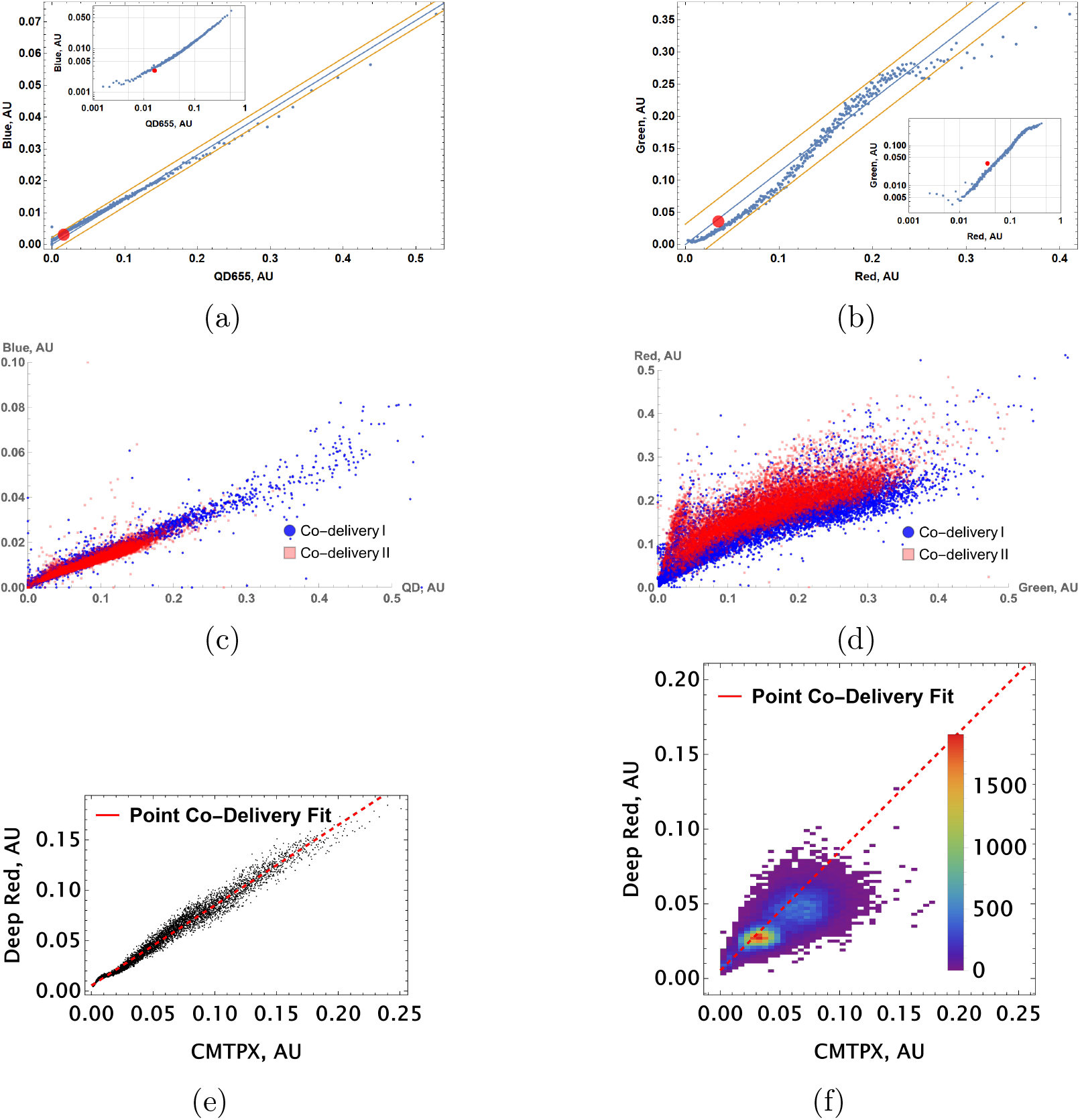
Coarse grained scatter plots for pairs of point co-delivered drugs are shown for (a) {Blue, QD_655_} and (b) {Green, Red} dyes. Log-log scale of dye intensities are shown in insets. Here, the large, red-filled circle on each of the plots corresponds to average values of dye intensities for the homogeneous delivery method. The data from two different point injection experiments are compared for pairs {Blue, QD_655_} and {Green, Red} in (c) and (d). In these two experiments, the locations of point injections were altered to demonstrate the independence of co-delivered dyes parameterized from the geometric setting. (e, f) Point co-delivery of CellTracker dye pairs (CMTPX and Deep Red) in fixed cells: (e) coarse-grained intensity data and its linear regression fit (f) the combined scatter plot of single-cell intensity data for two separate experiments in which dyes were delivered at homogeneous concentrations. Here, we used the same molar ratio of dyes (1 : 1) as in the point injection delivery, but delivered dye combinations homogeneously at concentrations of either 5 *μM* or 10 *μM*. The bi-modality is due to the separation of individual distributions of single-cell intensity data for each of these experiments.

We also observed a high degree of linear correlation in the uptake of co-delivered dyes in fixed cells (Figure 3e). As in the case of live cells, homogeneous dye delivery to fixed cells can be predicted well by linear regression obtained from the point co-delivery data (cf., Figure 3f). The correlation in uptake (*ρ* = 0.79) for point co-delivery in fixed (suspension) cells is comparable to the correlation observed in experiments with live cells (*ρ* ≥ 0.8).

To go beyond the “toy” system of conjugation-free co-delivery in which we treat one of the diester dyes or quantum dot (QD) nanoparticles as a drug, we co-delivered an authentic drug, auranofin, and a conjugation-free tag (Orange CellTracker dye) to human pulmonary artery endothelial cells (HPAEC). Auranofin is a thioredoxin reductase (TRR) inhibitor that has been shown to increase substantially intracellular H_2_O_2_ levels. We utilized HyPer7, a genetically encoded fluorescent probe for H_2_O_2_ detection, as a reporter for the drug’s effect on cell phenotype [8]. The co-delivery of auranofin and conjugation-free dye was performed by point injection. The results of the experiment are shown in Figure 4, where we quantified the fluorescence of the barcode and the H_2_O_2_ sensor’s ratiometric reading from the imaging data using a coarse-grained approach.

**Figure 4:**
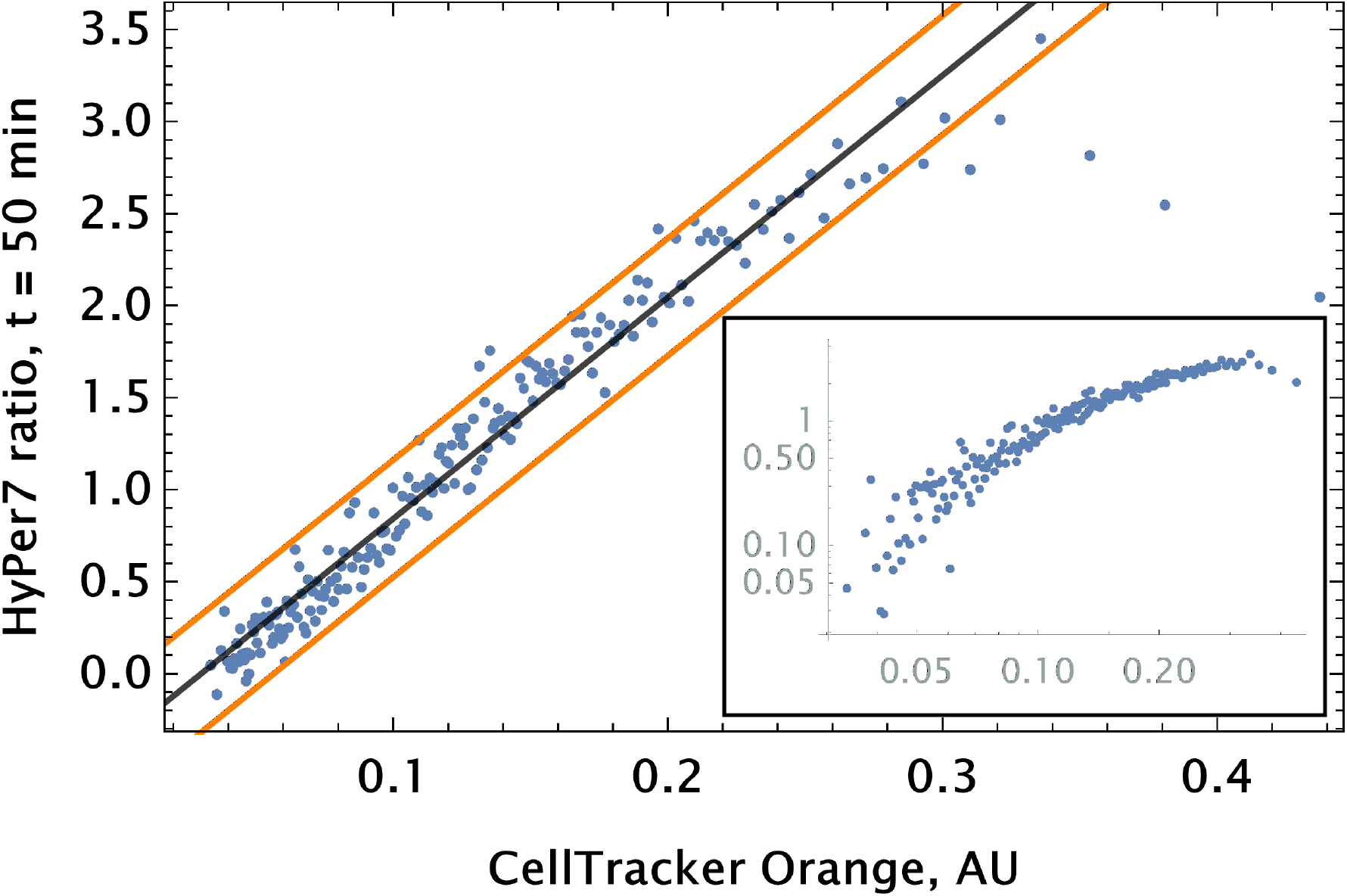
Auranofin conjugation-free barcoding in HPAEC: fluorescence signal of barcode (CellTracker Orange) and drug action proxy HyPer7 sensor (H_2_O_2_ fluorescent detector) were imaged by confocal microscopy. Quantification of the relationship between drug and dye uptake was performed using coarse grained analysis. The linear model fit for the HyPer ratio and 95% confidence intervals are shown by black and orange lines, respectively. The same scatter data are shown on log-log scale in the inset to demonstrate the broad log-linear range (~ 100 fold) of tag and drug correlation.

We observed a significant degree of correlation (*ρ* ~ 0.5) between the barcode tag for the drug and the drug effect as assessed by corresponding fluorescence signals (Figure 4). The observed relationship was largely linear with saturation in the ratiometric signal for H_2_O_2_ detection in cells with high auranofin uptake levels.

### Effect of siRNA Combinations on Gene Expression

Our benchmark system (dye combinations) was designed to demonstrate random drug mixing capabilities. We next considered a biological system with a detectable drug mixing effect that could be monitored. To do so, we chose drugs that are direct binding partners of a common, measurable target. We also wanted this system to be as portable as possible, i.e., be adaptable to different cell types, culture conditions, and, eventually, to *in vitro* testing.

We implemented this system as follows: The drugs’ target is the exogenous green fluorescent protein (eGFP) gene (d2EGFP, a short-lived variant, cf., Methods). The drugs are Small Interfering RNA (siRNA) duplexes targeting different regions of the GFP transcript.

We designed two classes of siRNA: (i) typical 21 nucleotide-long siRNA molecules targeting complementary binding sequences on eGFP mRNA; and (ii) “msiRNA,” i.e., siRNA with mismatch(es) in its targeting binding sequence. The msiRNA guide strand may still bind target transcripts (and, hence, interfere with target gene expression) via partial sequence complementarity by a mechanism closely mirroring microRNA (miRNA) silencing.

By analogy with endogenous miRNA in eukaryotic cells, one can expect weaker target protein repression by msiRNA compared to siRNA [9]. [Naturally occurring miRNAs can typically achieve significant repression of the target gene only by utilizing multiple/tandem binding sites on the target mRNA.]

We created more diversity in the drugs’ action by introducing two different classes of siRNAs targeting different sequence locations: (i) well separated and (ii) overlapping. The motivation behind this design followed theoretical considerations.

For low drug concentrations, we expected additive drug interaction regardless of the regions’ proximity since there is an abundance of target copy numbers to which siRNA molecules can bind. At high drug concentrations, we expected different drug interaction patterns that depend on target site locations. Different siRNA molecules targeting overlapping regions may, for example, antagonize each other due to binding competition.

For siRNAs targeting non-overlapping regions, one can expect additive or, perhaps, synergistic drug interactions owing to accelerated mRNA degradation and modulated translation rates in the presence of multiple RISC complexes bound to the mRNA. The siRNA design principles, specific sequences, and corresponding target region locations are shown in Figures 5a and 5b (cf. Methods for the details).

**Figure 5:**
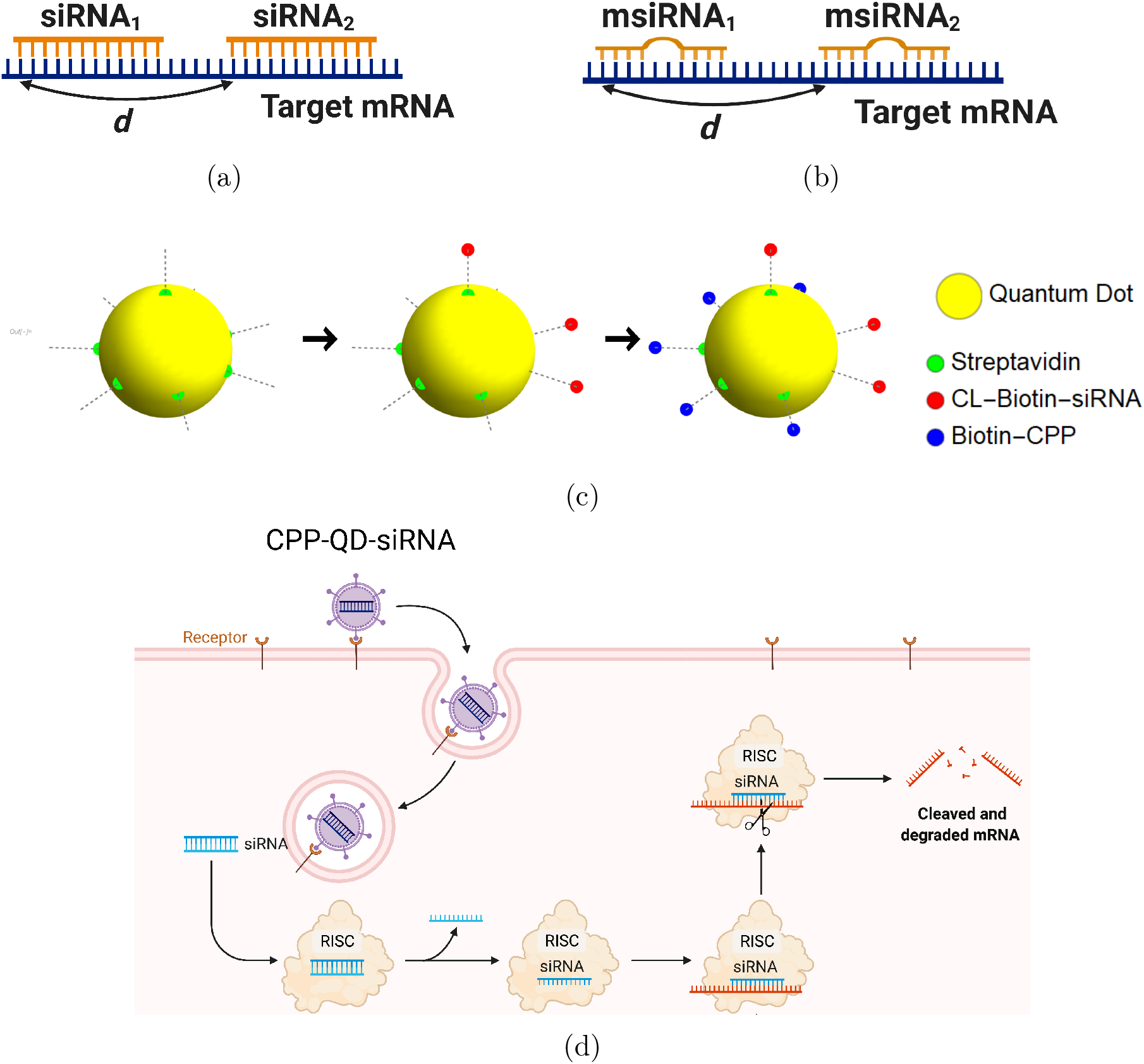
siRNA design: Different GFP messenger sites in open frame are targeted by either (a) fully complementary siRNA or (b) partially complementary msiRNA (b) small interfering RNA molecules of length 21 nucleotides. The distance *d* in nucleotide units is measured between the 3’ ends of targeting sites. The siRNA are non-overlapping if d ≫*22nt* and overlapping otherwise. Note that pairs {siRNA_*i*_, msiRNA_*i*_} completely overlap, d = 0. Each siRNA is conjugated to biotin molecules via a cleavable (disulfide) linker (CL). (b) The siRNA are delivered into cells by Quantum Dot nanoparticles (QD). The QD are functionalized with streptavidin molecules (5-10 per QD nanoparticle) which are used to load siRNA and biotinylated cell penetrating peptides (CPP) guiding and anchoring the fluorescent nanoparticles into the cells. The siRNA sense strand was designed with a cleavable linker (CL) at 5’ end. The linker region contains a disulfide bond, which releases the siRNA within the reducing cytosolic environment.

Cell-penetrating peptide-conjugated, nanoparticles, Quantum Dots (QD), were used both for drug delivery and barcoding (Figures 5c and 5d). We delivered QD-conjugated siRNAs targeting GFP mRNA in a pairwise fashion. In these experiments, we point-injected all possible combinations of three different siRNAs: siRNA_1_, siRNA_2_, and msiRNA_1_. As expected, point injection of a drug conjugated and delivered in this way introduces a broad range of drug uptake across the sample (Figure S10), as compared to the homogeneous random mixing approach (Figure S9b), and suppresses the proportionality bias in drug combinations.

The siRNA_1_ and siRNA_2_ species have perfect complementarity to non-overlapping regions of mRNA. The msiRNA_1_ was designed to bind to the same region of mRNA as siRNA_1_ but has a mismatched nucleotide pair within the targeting sequence. The siRNAs were barcoded using two optically different QD nanocarriers, QD_605_ and QD_655_. To demonstrate that the drug combination effect is independent of the tag, we exchanged the tags for each siRNA mixing experiment we performed.

Inherited randomness in cell behavior is not only responsible for variable drug uptake, but also all but guarantees variability in response to drug treatment by individual cells. Even under conditions of identical uptake of a drug and the same exposure time, one expects heterogeneity in individual cell response/phenotype. To overcome this source of heterogeneity, we perform averaging over many cells within each sub-populations characterized by a specific combination of tags (and hence drugs). The number of cells in each subset must be large enough to suppress the variability in drug response ensured by the Law of Large Numbers.

The results of siRNA combinations on target gene expression are shown in Figure 6. Here, we partitioned the QD tag intensities into the 2D bins spanning effective parametric space. GFP intensity was averaged across a subpopulation of cells in each bin. Since the QDs at high concentrations lead to cytotoxicity, GFP intensity for each condition was normalized to the control sample in which we point-injected *QD*_605_ and *QD*_655_ nanoparticles not conjugated to siRNA. (This strategy is equivalent to normalization of repression levels using scrambled siRNA samples as controls.) We used a machine learning algorithm to predict the normalization factor for QD combinations using this control sample (see Supporting Information for details).

**Figure 6:**
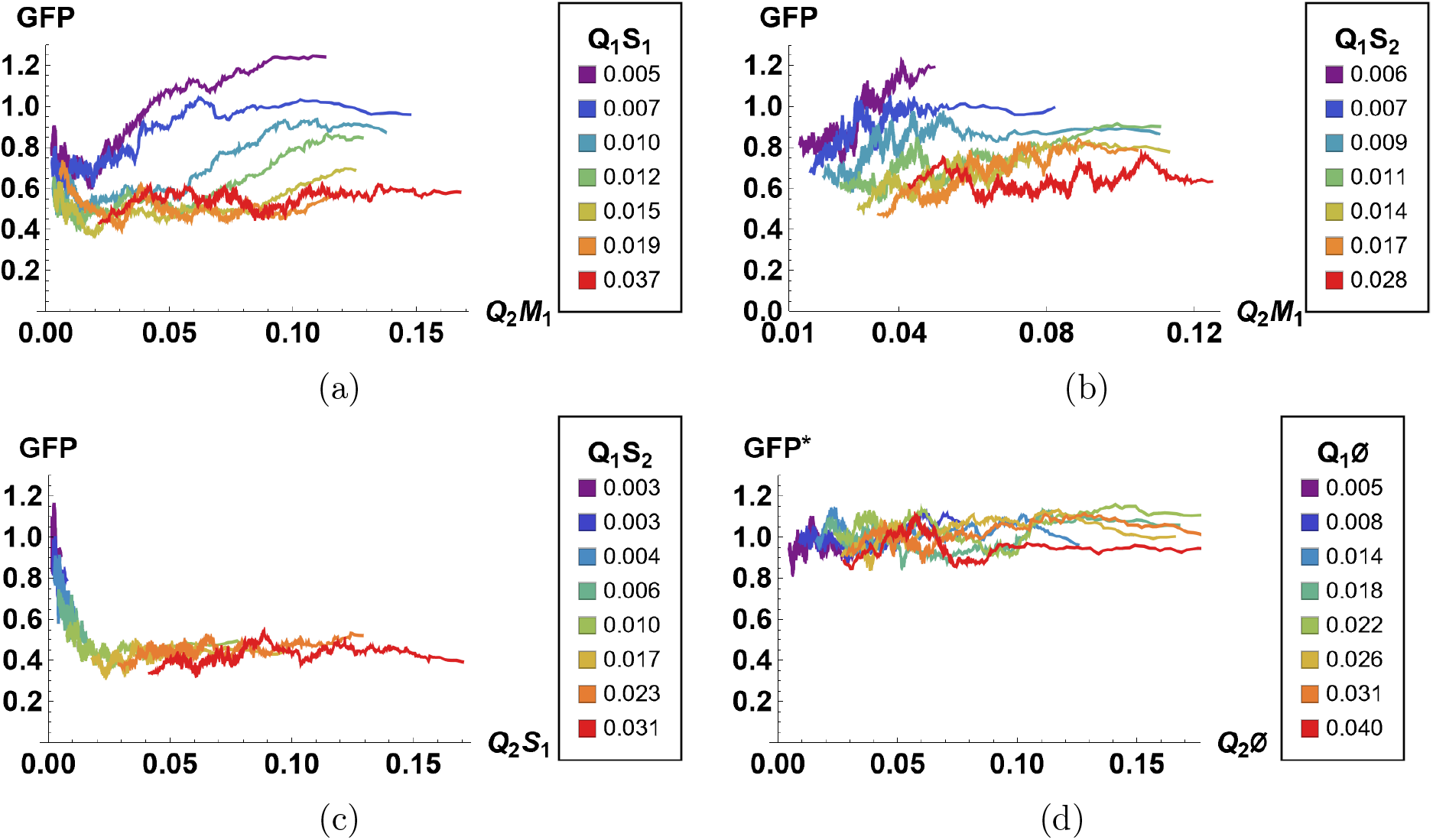
The effect of siRNA combinations on target gene (GFP) expression. Three different siRNAs were delivered to cells by optically distinguishable QD nanocarriers (*Q*_1_ and *Q*_2_) in pair-wise fashion. Two siRNAs, siRNA_1_(*S*_1_) and siRNA_2_(*S*_2_), were designed to bind non-overlapping sequences of the mRNA target in completely complementary fashion. The third siRNA, msiRNA_1_(*M*_1_), has almost identical primary structure as siRNA_1_ with a single nucleotide mutation close to the seed region. (a, b) Relief of GFP repression mediated by msiRNA_1_ is shown for different concentrations of the repressors siRNA_1_ and siRNA_2_, respectively. (c) Combination of non-overlapping repressors siRNA_1_ and siRNA_2_ results in additive/synergistic repression of target GFP expression. (d) Control sample of siRNA free QD combinations was used to account for QD toxicity for both carriers using a machine learning algorithm, and its predicative accuracy is shown here for different concentrations of *Q*_1_ and *Q*_2_.

At high concentrations, the “mismatched” siRNA, msiRNA_1_ (with incomplete complementarity in its target binding sequence), acts as an inhibitor for the strong repressors siRNA_1_ and siRNA_2_, as shown in Figures 6a and 6b. This antagonistic effect is more pronounced in the siRNA_1_-msiRNA_1_ combination as compared to the siRNA_2_-msiRNA_1_ combination.

The outcome of pairwise combinations of the non-overlapping strong repressors siRNA_1_ and siRNA_2_ suggests additive or synergistic interaction for these drugs (Figure 6c). The bias in the titration range of the siRNA concentrations in this experiment is due to limitation of effective mixing conditions (see Supporting Information for details). The predictive accuracy of the machine learning algorithm for GFP normalization (due to toxicity of QDs) is shown in Figure 6d for a broad range of mixing conditions.

Here, we, once again, employed the tiled image renormalization approach, which is especially well suited for point injection data analysis because local gradients in drug concentrations persist over scales sized larger than that of a single cell. Since tiles in the image relate to cells in the same spatial proximity, tiled renormalization is equivalent to concentration(s) gating by the local microenvironment.

The results of the pairwise drug combinations in Figure 6 demonstrate both additive and antagonistic effects of the siRNA combinations. We note that direct binding competition is not the only source of possible antagonistic siRNAs interaction. The repression by siRNA is mediated by the RNA-induced Silencing Complex (RISC). At the high intracellular concentration of siRNA duplexes, one may expect competition for RISC integration by different siRNA species due to the limited pool of RISC in the cell. This type of competition may result in antagonistic siRNA interactions, regardless of their target [10, 11].

### Random mixing and response quantification of three and four drugs

To demonstrate that our approach can be applied to combinations of more than just a pairs of drugs, we considered the following application. We tested drugs commonly used in combination chemotherapy regimens to treat different forms of cancer, such as cyclophosphamide (CP), docetaxel (TXT), 5-fluorouracil (5-FU), and cisplatin (CIS). The desired phenotype in this application is cell death, which can be assessed by utilizing fluorescent biomarkers, such as Annexin-V stain.

Alternatively, death phenotype can be assessed by monitoring cell detachment and/or changes in cell morphology, which may be resolved by bright field/phase contrast microscopy. Cell shrinkage due to dehydration and cell debris can also be detected by a decrease in intensity of the forward light scatter (FSC) signal in flow cytometry measurements [12]. Unlike annexin-V staining, these cell death detection methods do not require, in principle, additional fluorescence biomarker(s) and, hence, do not reduce the necessary bandwidth required to resolve quantitation of drug combinations in individual cells.

We used the HeLa cell line as a benchmark system to evaluate the effect(s) of drug combinations in individual cells. We utilized either time resolved confocal microscopy, flow cytometry, or a hybrid of both detection methods in our experiments. In all experiments reported below, cells were treated with drugs for a short duration (1/2 hour). Hence, drug concentrations were chosen to be significantly higher than their corresponding typical *IC*_50_ values. (In typical experiments, drug(s) incubation is 48 hours, roughly 100 times longer than in our setting.) In order to create a negative drug control, cisplatin was inactivated by DMSO [13].

We note that our goal was to demonstrate the technical feasibility of our stochastic mixing approach. A more practical, generic application (in terms of both an expanded list of drugs and of cell lines) and biological data interpretation are beyond the scope of this paper.

### Flow cytometry measurements of cell death phenotype, three-drug combinations

In order to assess drug combinations leading to cell death using flow cytometry, we harvested detached cells and debris in the sample media at different time points and analyzed the fluorescence intensities of the specific drug tags, which correspond to *effective* drug combinations. At the final time point of the experiment, we harvested the remaining attached cells and analyzed the fluorescence intensities of the specific drug tags similarly, which correspond to *ineffective* drug combinations.

We stained cells homogeneously with the same auxiliary dye (Deep Red CellTracker dye) not conjugated to any drug. We used an identical concentration of this dye in all samples, including controls. The CIS, CP, and TXT drugs were labeled with blue, green, and orange CellTracker dyes respectively. All CellTracker fluorescence probes (except for Deep Red) contain a chloromethyl or bromomethyl group that reacts with thiol groups, utilizing a glutathione-S-transferase–mediated reaction.

In most cells, glutathione levels are high (*mM* range) and glutathione transferase is ubiquitous. The CellTracker Deep Red probe contains a succinimidyl ester functionality, which reacts with amine groups present on proteins, and, hence, is also ubiquitous in cellular cytoplasm. The ratio *R_i_* of any CellTracker dye *i* signal to Deep Red intensity can be thought as a proxy to cellular concentration of the CellTracker and its cognate drug, since the Deep Red signal is proportional to cell volume.

We assumed that for sufficiently large cell fragments this normalization also holds, which allows an estimation of the drug concentrations in both intact cells and dead cells (debris). We note that due to stochastic uptake of any dye including an auxiliary dye, this normalization is only meaningful after averaging over a significant number of detection events required to suppress the signal noise.

In order to convert the ratio *R_i_* to absolute drug concentration, we calculated 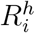 for a homogeneously delivered (with mixing) drug sample where the drug concentrations are known exactly. The volume concentration of drugs in these homogeneously delivered controls were {1, 1, 0.1 *μM*} for NC, CP, and TXT respectively.

The factor 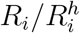 calculated for individual cell is a proxy for the volume concentration of the drug *i* compared to homogeneously delivered drug control. Given the ubiquitous nature of all CellTracker dye targets, one expects that this estimate will hold even after cell division. The absolute fluorescent signal from the mother cell will decrease due to dilution by cell division, but the ratio of homogeneously distributed targets should persist.

To demonstrate that this expectation holds, we considered dynamic changes in fluorescence signal intensity in the controls of homogeneously delivered drugs. The population average factors, *R_i_* for *i* = {NC, CP, TXT} are shown in Figure 7a at different time points.

**Figure 7:**
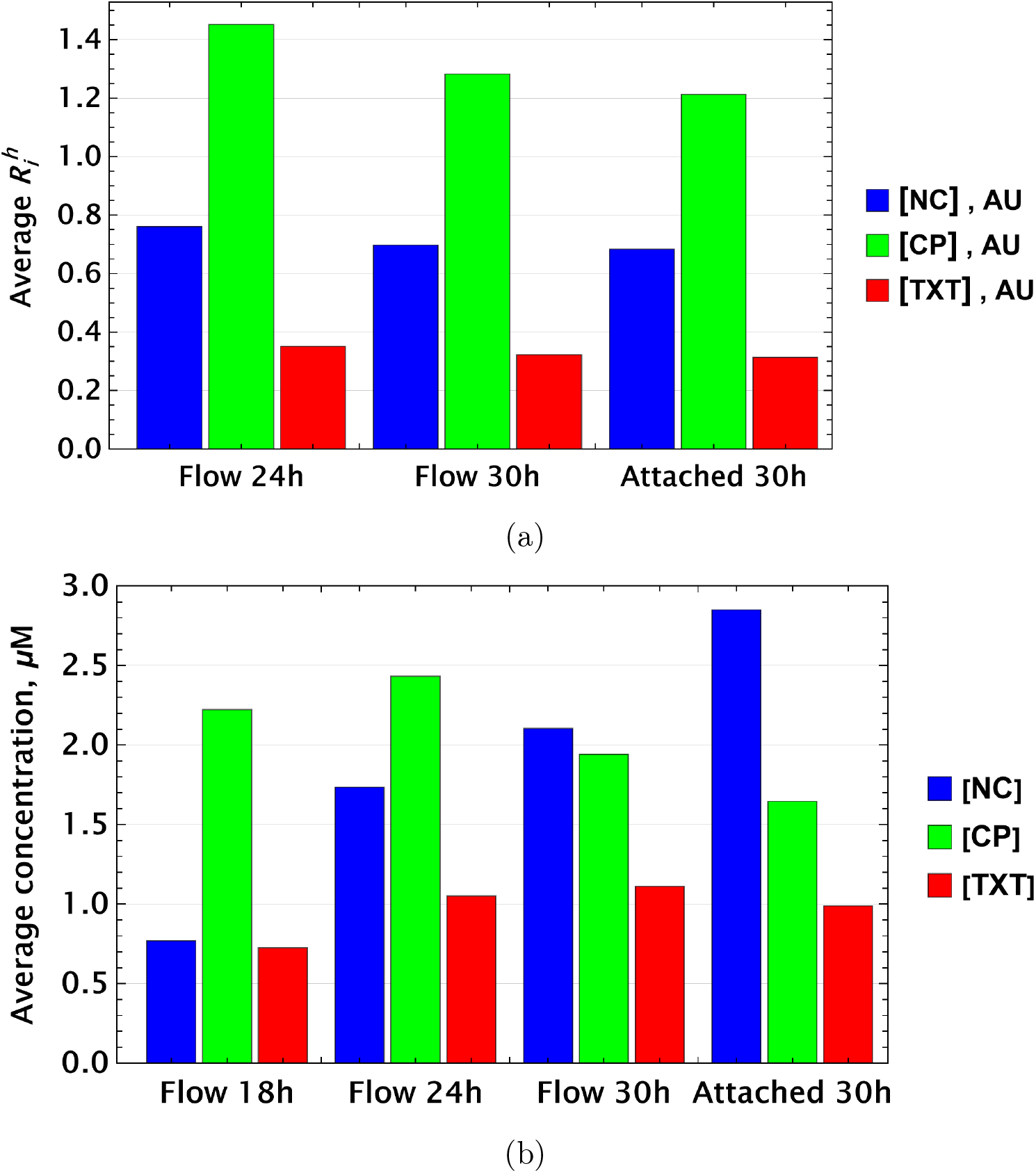
(a) Population average of {[*NC*], [*CP*], [*TXT*]} drug concentrations in homogeneous delivery (thoroughly mixed) experiments, shown in blue, green, and red, respectively. Drugs’ concentrations are measured with respect to an auxiliary or marker dye. The first two samples correspond to detached cells harvested from the sample media at different time points and the last sample corresponds to surviving attached cells harvested at the final time point. (b) The same analysis as in (a) applied to non-homogeneous (i.e., gradient or point-delivery methods) drug delivery experiments. Drugs’ absolute concentrations were determined using a linear normalization method based on intensities from homogeneously delivered samples. The first three samples correspond to detached cells harvested from the sample media at different time points, and the last sample corresponds to surviving attached cells harvested at the final time point.

Small time-dependent modulation of 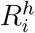 values are noted and are a consequence of the stochastic nature of drug or dye uptake, a process that ensures the existence of cells with a distribution of drug concentrations, including importantly some that take up less drug(s) than the cell average. The corresponding *R_i_* values for these cells is smaller compared to cells that received higher drug dosage. Hence, this low *R_i_* subpopulation is more likely to survive longer. Since this dynamic drift is rather small, however, we will neglect it in further analysis.

For samples created by gradient drug mixing, one naively expects qualitatively the following picture for the population average of drug concentrations: the concentrations of effective and ineffective drugs should decrease and increase, respectively, with time. This expectation derives from the fact that the subpopulation of cells that received ineffective drug concentrations or drug combinations becomes increasingly dominant over time.

This qualitative picture (evaluated on a whole cell population level) holds only if drugs act independently in individual cells. Drug pharmacodynamics may also be non-uniform over time. We observed these behaviors in the gradient-derived mixed sample data shown in Figure 7b.

The dynamics of inactivated cisplatin (labeled NC in fig. 7b), indeed, displays the time dependence expected from an ineffective drug (concentration). It is much more difficult to interpret the pharmacodynamics of CP and TXT because the time dependence is not monotonic. Drug-drug interactions, drug effect on cell cycle, and other time-dependent drug actions can all play a significant role in overall cell population behavior.

To visualize the interplay of drugs on individual cells, we first reduced data dimensionality by discarding information about the level of [NC] in cells (assuming DMSO inactivated cisplatin completely). Under these circumstances, one can plot the probability distribution, *P*(*x,y*), to find cells with different drug combinations {*x* = [CP], *y* = [*TXT*]} at different time points, as shown in Figure 8a. Two dimensional projections of this histogram for each individual time point are shown in Figure 8b.

**Figure 8:**
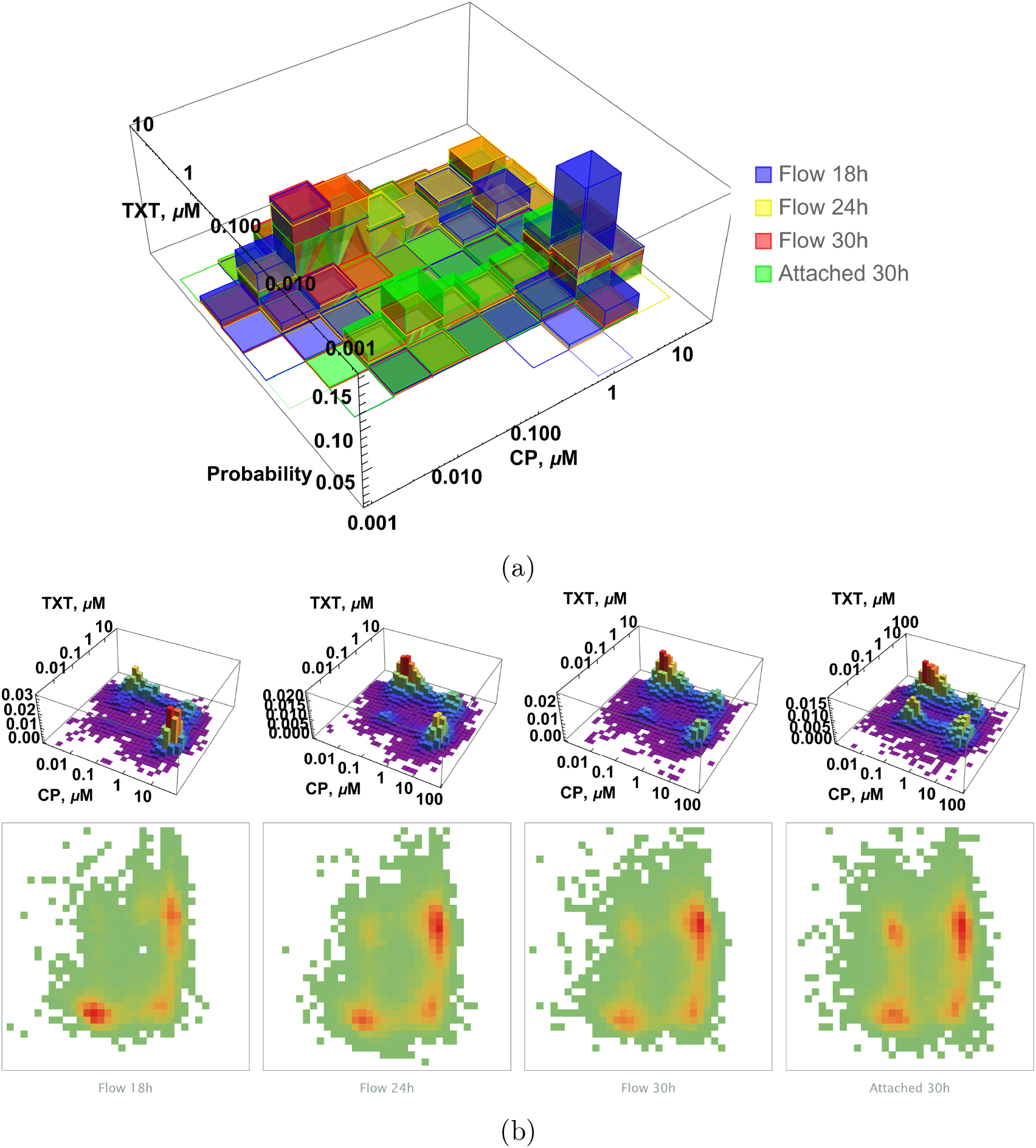
(a) Histogram of {*CP, TXT*} pairwise drug distribution in intact cells. Different colors correspond to detached cells harvested from the sample media at different time points (shown in blue, yellow, and red, respectively), and surviving attached cells harvested at the final time point (shown in green). (b) Time resolved changes in drug content of dead (first three samples) and live (last sample) cells collected. The density plots represent cell count as a function of drug concentrations (log scale) at different time points.

Our goal was to demonstrate technical capability of stochastic mixing approach. More practical application (in terms of both list of drugs and cell lines) and data interpretation are beyond the scope of this paper.

### Microscopy measurements of cell death phenotype, four-drug combinations

For microscopy-based measurements and image analysis, we used the point injection method to create local drug gradients across the sample of confluent HeLa cells on a glass slide. Followed by the initial imaging of the drugs’ distributions across the sample, we monitored cell detachment (and changes in morphology, e.g., shrinkage) at different locations on the slide at different time points. The estimated intracellular concentrations of all chemotherapy drugs used were assessed using the non-conjugated fluorescent tag (dye) method described above.

The image analysis allows one to correlate the initial drug combination concentrations in each locale on the slide and subsequent death phenotype in the same position at different time points. In principle, cell migration between locales can complicate the analysis, but we used confluent seeding conditions to minimize this effect.

The outcome of a typical microscopy experiment is shown in Figure 9. Here, the initial distribution of each drug across the sample is shown on the first panel in Figure 9 using four different colors (blue, green, red, deep red) corresponding to the fluorescent label for each drug. Local cell density at different time points is shown in Figure 9, panels 2-4, using an inverse rainbow color scheme (where blue and red correspond to fully confluent cells and no cells, respectively).

**Figure 9:**
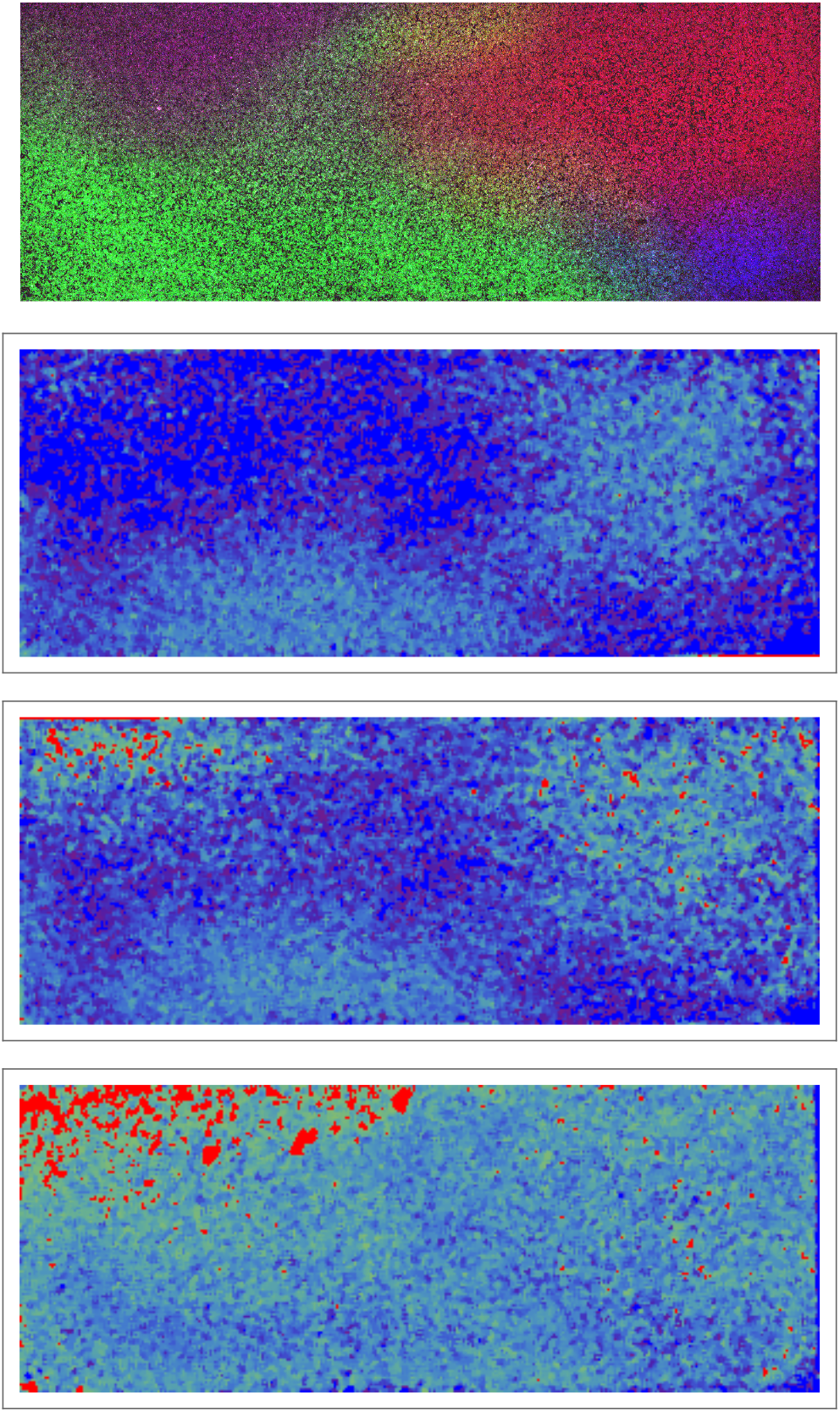
Time resolved measurement of attached cell density. Top row: initial spatial distribution of drug concentrations of NC, CP, TXT, 5-FU is shown in blue, green, red, and dark red respectively. Rows 2, 3, 4: calculated cell local cell density at time points 18, 24, 30 hours, respectively. Local cell density is determined by the fraction of cell-covered area within a square of 10^4^ *μ*^2^. High and low cell confluency correspond to fractional range from 1 (blue) and 0 (red) respectively.

We used a coarse-grain approach for image analysis, averaging fluorescence intensities over an area of ~ 10^4^*μm*^2^ at each locale. Cell density was assessed by cell segmentation using bright field imaging data and the calculation of the area fraction covered by cells in each coarse-grain locale. The absolute drug concentration can be determined using the homogeneous delivery method discussed above.

## Discussion

Biologists often treat noise in samples as a nuisance that must be minimized, an important consideration for typical experimental measurements. Here, we rely on engineered local heterogeneity in drug uptake as a tool to encode drug treatment regimens and to predict the macroscopic response to drug perturbations. This approach was inspired by the celebrated fluctuation–dissipation theorem in statistical physics [14] that predicts the response of a system to external perturbations based on the underlying properties of noise in the system at equilibrium.

Rather than testing thousands of samples, as little as a single cell culture sample is required to perform the search for optimal drug combination using our approach. Applications of our method are not limited to the identification of optimal drug concentrations, as the approach is applicable to any quantifiable cell phenotype. The method is readily generalizable to a much broader set of problems involving complex biochemical or molecular perturbations applied *in vitro* and *in vivo*. Examples of these perturbations include combinations of signaling molecules, optimization of drug delivery strategies, lineage differentiation identification, stem cell manipulations and tracking, etc.

The best possible random drug mixing is achieved in cells in suspension. Suspension cells can be easily mixed (i.e., cell locations can be randomly re-distributed), and local drug gradients can be applied in a statistically independent serial fashion. By contrast, since the locations of adherent cells in the sample are fixed, we exclusively rely on randomness of the drug distribution in the media by means of forced convection, which is challenging to control. Hence, further theoretical and experimental work is required to improve the stochastic drug mixing range and its smoothness for adherent cells. A possible way to improve mixing range (and save reagents even further) can be utilization of recently developed “Dye Drop” method [15].

We would like to point out an important limitation of our stochastic drug mixing method (at least in its current state). Throughout this paper, we tacitly assumed that both drugs and their labels are retained by the cells and are not re-distributed from cell to cell after the washing step. This is certainly the case for siRNA drugs delivered via nanoparticles and CellTracker dyes in our applications, but this assumption does not necessarily hold in general.

Drugs can, in principle, leak from the cells into the culture media after the washing step and, eventually, be re-distributed homogeneously across the entire cell population. This behavior will affect the “area under the curve” for individual cells after prolonged incubation periods, especially cells that had initially very low (or no) drug exposure. We assumed that this effect is small and did not study the kinetics or the extent of this “secondary” drug exposure here. (Multiple washing steps were performed to mitigate this effect.) More careful analysis and “compensation” is required for drugs that are not retained by the cells.

Another possible drawback of our method is a limit on maximal drug concentration due to drug solubility and its solvent (e.g., DMSO) toxicity. Since we utilize short drug exposure times, high drug concentrations may be required to test a broad range of conditions. Typically, low concentrations of individual drugs are desired in combination therapies and, hence, our method can be implemented even for drugs with low solubility.

We expect possible adaptation of the point injection method to facilitate random mixing *in vivo*. The spatial distances between injection sites, geometry, needle gauge size, injection volume, and the specific measurement protocol (readout) will, of course, all need to be established and optimized for a particular application.

## Materials and Methods

### Drug labeling and delivery

One barcoding option is to label the drug directly with a fluorescent tag either by chemical modification of the drug or other irreversible binding methods. Here, we demonstrate the application of our method for irreversibly tagged drugs using cell-retained dyes as a benchmark drug model. The cell-permeable diester dye, carboxy fluorescein succinimidyl ester, and its derivatives are an example of this type of drug.

We used multiple different dyes to perform the multiplexing experiments: CellTracker dyes (blue, green, orange, CMTPX red, and deep red) and CellTracer dyes [violet, carboxy fluorescein succinimidyl ester (CFSE), yellow, red, and far-red]. The CellTracker probes have been designed to pass freely through cell membranes, and once inside the cell they are converted into impermeable fluorescent products. The probes are transferred to daughter cells, but are not transferred to adjacent cells in a population. The CellTracker fluorescence probes contain chloromethyl/bromomethyl or succinimidyl ester reactive groups that react with ubiquitous intracellular thiol or amine groups, respectively.

More flexible barcoding can be achieved by utilizing a tag bound to a drug by a cleavable linker. The tag can be designed to be retained by the cell, with the delivered cellular drug amount proportional to the tag’s fluorescence signal. We used nanoparticle drug delivery to apply this labeling method, namely, cell-penetrating peptide-conjugated quantum dots (QD) were used both for drug delivery and barcoding. The fluorescent reporter (QD) is retained within the cell upon entry and the drug, siRNA in our case, is released locally into the cytosol via cleavage of a disulfide bond in the cytosolic reducing environment.

Finally, we utilized the point injection method to co-deliver locally both drug and conjugation-free fluorescent dye retained by the cells. Here, we rely on the similarity of temporal concentration profiles of drug and dye within cell culture. Drug (and dye) uptake is governed by transport and diffusive processes, as well as molecular interactions with its targets, both specific and non-specific [7].

In many cases, strong, specific drug binding to the target leads to fast reaction rates which, in turn, result in the uptake process being diffusion/transport-limited. In this situation, media concentration profiles created by the *initial* dispersion drive intracellular drug and dye uptake. We pre-mixed reagents used for co-delivery in a small volume that is injected into a much larger sample volume. Hence, point co-delivery results in similar local concentration profiles of drug and dye, at least for short incubation periods.

We note that to achieve one-to-one (monotonic) correspondence between total cellular drug uptake and its conjugation-free tag, it is important to ensure non-saturation labeling and detection conditions. If the tag fluorescence intensity saturates beyond a certain exposure concentration (or time), the drug total uptake prediction will be inaccurate. Most of the dyes we used (CellTrace and CellTracker) have abundant intracellular targets and saturate only at very high extra-cellular concentrations (> 1 *mM*). Therefore, the saturation in tag signal may be caused by microscopy detection limits (for either too low or too high fluorescent signals at the given laser power and sensitivity of the photomultiplier settings). Careful adjustment of these parameters is required to ensure a maximal dynamic detection range.

### Effective drug combinations by stochastic mixing

Stochastic mixing occurs naturally due to randomness in the drug(s) uptake process by individual cells. We enhance the random process of drugs’ uptake by creating transient drug gradients across the sample.

In the simplest possible implementation of *determinitic* gradient mixing, one can create a drug *A* gradient along the sample’s x-axis and a drug *B* gradient along its y-axis, generating (locally) drug *A* and drug *B* combinations in a continuous fashion. This approach, however, requires microfluidic devices that would maintain stable drug gradients, which can be technically challenging [16, 17]. Moreover, the implementation of multiple (i.e., > 2) drug gradients *ex vivo* is probably not feasible on a small spatial scale. It is also not obvious as to how to prepare efficient combinations of three or more drugs using this approach.

Here, we used a much simpler approach utilizing local point injection to create spatiotemporal drug gradients in (mammalian) cell culture. We do not homogeneously distribute drugs in the media but, rather, inject them locally into cell culture in separate, discrete locations. Point injection of multiple drugs ensures chaotic/stochastic mixing patterns across sample. The illustration of the stochastic mixing method implemented on a much larger scale is illustrated by aesthetic visualization below:

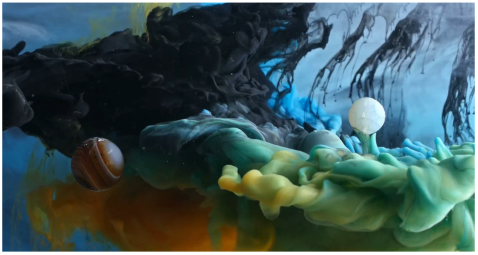
Ink In Motion by Macro Room, https://www.youtube.com/embed/ICxC5ekWnUc.

One can think of transient drug gradients as local, short incubation processes that facilitate uptake heterogeneity across the sample. The duration(s) of these local uptake processes are governed by diffusion, convection, and the drug-target reaction kinetics in cell culture.

Steps involved in the point injection method compared to homogeneous delivery are shown in Figure 10a and Figure 10b, respectively. Cells in suspension culture were placed in a cuvette and a dye *A* (Red Dye in the example shown in Figure 10b) was injected locally (just beneath the meniscus). The volume of the point injection must be small compared to the cell culture volume in the cuvette (we used ~ 1 : 20 volume ratio). (N.B., the graphic illustration in Figure 10b is an idealization because the gradient created by point delivery is not spatially linear.)

**Figure 10:**
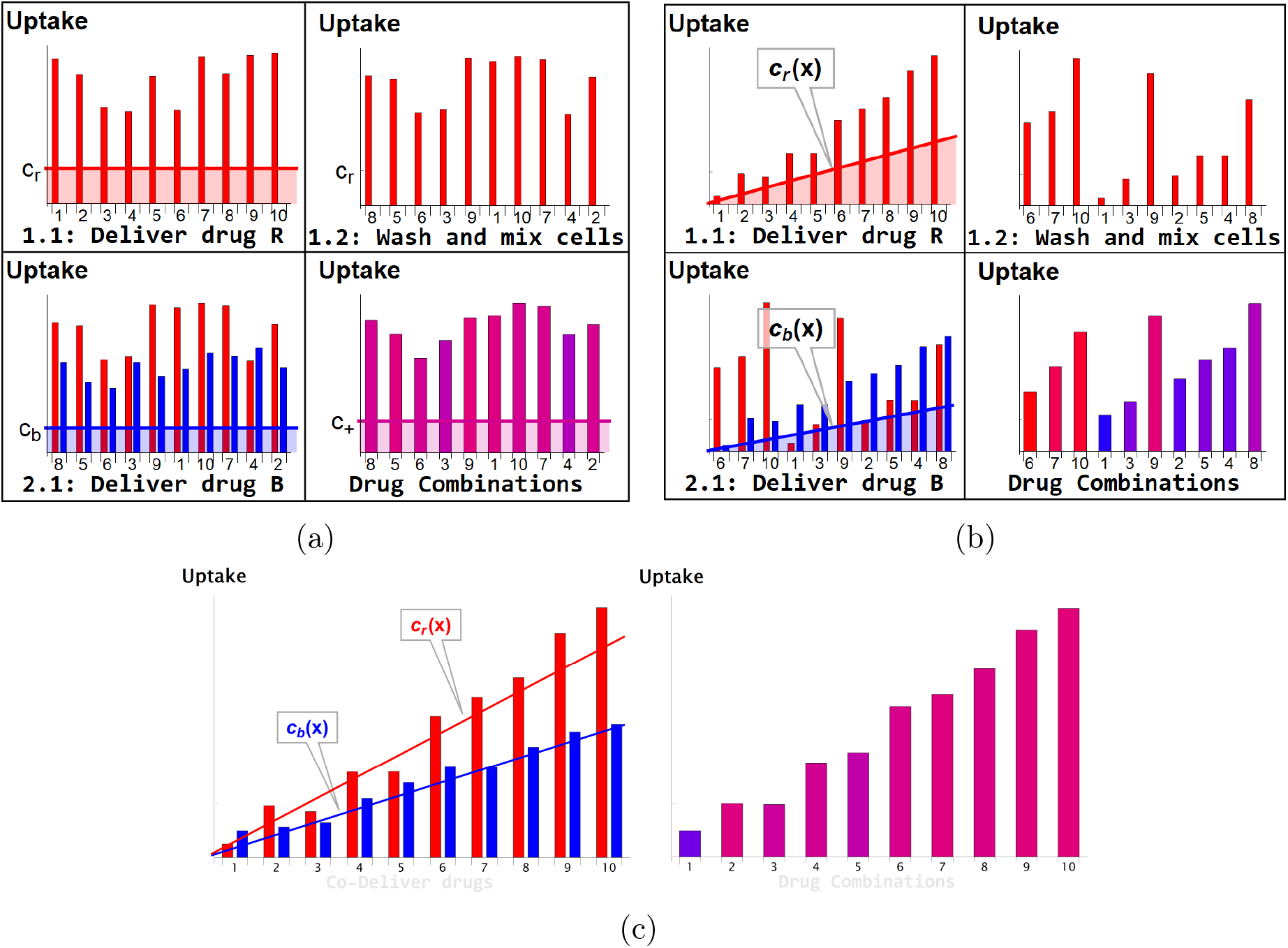
(a) and (b), Graphic idealized illustrations of dye/drug delivery are shown for homogeneous and point injection methods, respectively. Here, x-axis values correspond to the *initial* position index of individual HeLa cells. Media dye concentration profiles are depicted by shaded regions on the left panel for both delivery methods. The y-axis values represent cellular dye uptake for each stage of staining (panels 1.1, 1.2, and 2.1). The final dye combinations are shown in the bottom right panels for both delivery methods (the bar color and the amplitude reflect the molar ratio and total uptake of dyes, respectively). Each wash step is mimicked by reshuffling the cell position index (cf., panels 1.1 and 1.2). Variation in dye uptake due to intrinsic cellular heterogeneity is illustrated for each of the delivery methods by imposing multiplicative noise. For simplicity, the noise in uptake is assumed to be independent for each dye. (c) Graphic illustration of the point co-delivery method.

The cultured cells suspended in the cuvette were left undisturbed during the short incubation process (10-30 minutes). The transient gradient of dye *A* results in higher dye uptake by cells located closer to the point injection site. The subsequent wash step ensures removal of remaining dye *A* and also mixing of differentially stained cells. The process is then repeated with another dye *B* (Blue Dye in the example shown in Figure 10b) in exactly the same fashion as with dye *A*.

As a result, a broad range of drug combinations can be achieved. Consider, for example, cells in the proximity of the dye *B* point injection site. These cells exhibit a wide range of dye *A* uptake and high uptake of dye *B*. By contrast, cells remote from the dye *B* point injection site are also characterized by a broad range of dye *A* uptake but low uptake of dye *B*. The conjugation-free barcoding method to co-deliver locally both drug and fluorescent dye retained by the cells is illustrated graphically in Figure 10c.

The local injection method was used to create stochastic drug combinations in adherent cell culture as well. (Cell mixing, naturally occurring in suspension cell culture, is not feasible in this case unless cells are detached and re-seeded, actions which require time and may interfere with drug response.) The local drug gradients across the single sample created using this method are driven mostly by initial drug dispersion (forced convection) during the injection if the incubation time is short (cf., Supporting Information for technical details on diffusion rate of small molecules).

The results shown in Figures 2 and 3 correspond to experiments performed with 90% confluent HeLa cells cultured in 4 wells with a chambered coverslip Ph+ (Ibidi). Point injections were performed pairwise: pair (i) CellTrace Violet and Qtracker 655 Cell Labeling Kit (referred to as Blue and QD655 dyes in the main text); pair (ii) CellTrace CFSE and CellTracker Red CMTPX (referred to as Green and Red dyes in the main text).

The total culture volume of each chamber was 750 *μl* and 30 *μl* were replaced with a pre-mixed pair of dyes injected at one of the chamber corners. The process was repeated with the second pair of pre-mixed dyes injected into another chamber corner. The manual injection rate is difficult to control but the total release time was ~ 1 sec. The incubation was performed at room temperature for ~ 20 min in the cell culture hood (with all possible precautions taken to avoid shaking the slide).

The graphic illustration in Figure 11 demonstrates this approach for a pair of drugs. Here, an idealized drug mixture distribution is depicted by color and intensity in Figure 11a. All fluctuations due to cell variability, convection, and non-independence in drug uptake are ignored in these illustrations for simplicity. Moreover, we assumed that transient drug uptake gradients are symmetric (radial) with respect to the point injection locations (left and right top corners of the rectangular slide). In reality, of course, all of these assumptions, including radial symmetry, are violated, resulting in much more random behavior, which is precisely what we wish to exploit to enhance the heterogeneity in drug uptake across the cell population.

**Figure 11:**
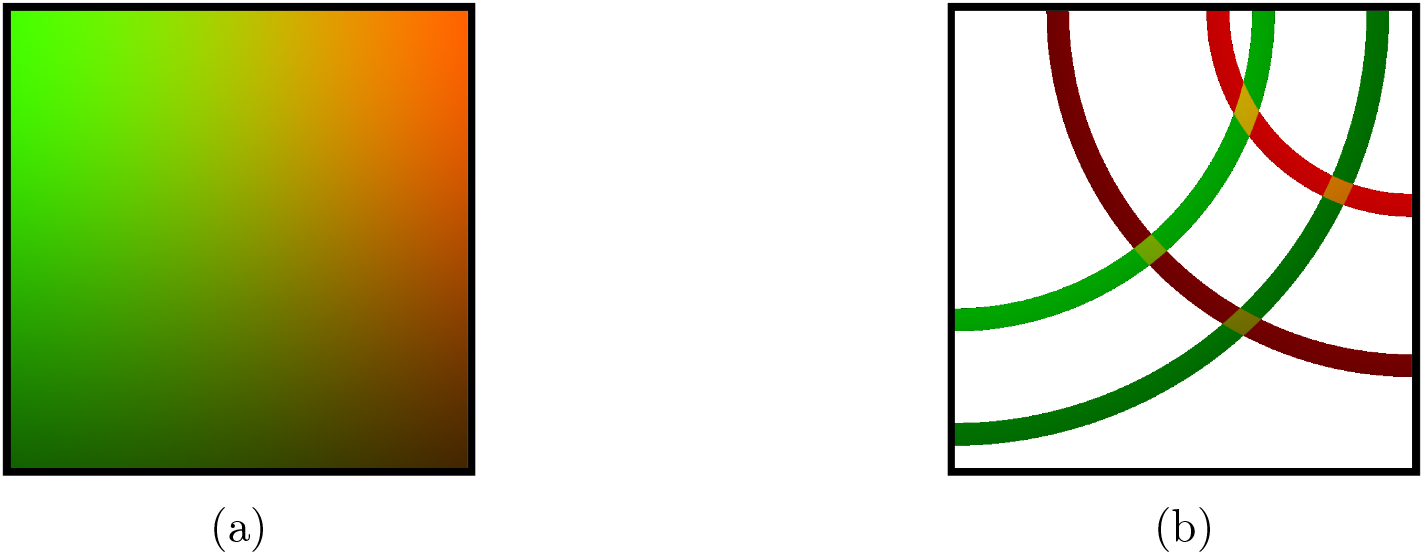
(a) Graphic illustration of the point injection method for adherent cells in culture. The idealized spatial distribution of two drugs (dyes) is shown, assuming they were delivered in the left and right upper corners (green and red color intensities correspond to drug uptake by cells). The fluctuations due to environmental factors and cell heterogeneity are ignored for simplicity, as well as the ripple effect due to finite well size. b) Discretization (binning) of uptake for each drug corresponds to spatial partitioning by concentric annuli of finite width around the point injection origin. The intersection of the annuli for two different drugs corresponds to unique drug combinations (four different conditions, or annulus intersections, are shown).

In Figure 11b, the origin of the distribution as a linear superposition of individual drug uptakes is shown, which is designed to illustrate that unique mixing conditions, even in this idealized case, depend on the geometry of the matrix and point injection locations. Discretization (binning) of uptake for each drug has a simple geometric interpretation in this idealized case, shown as regions of annular intersections in which colors are uniquely mixed. For a real system, this symmetry is violated due to processes such as convection, and we, therefore, must rely on the fluorescence intensities of the tags rather than geometry to partition the ensemble into bins; however, as in the case of the idealized system in Figure 11b, one expects a finite-sized local region on the slide where drug concentrations/uptakes can be assumed to be constant.

### Statistical processing of single cell data

In all single-cell experiments multidimensional intensity data were compiled using averaged fluorescence intensity measurements of each optical drug barcode for each cell. Additionally, the ensuing phenotype for each individual cell was recorded using the corresponding fluorescence signal in functional experiments (conjugation-free barcoding, auranofin, and siRNA combinations).

For experiments utilizing a single barcoded drug, we used coarse-grained modeling to recover the relationship between drug and tag as follows: all measured single-cell fluorescence intensity data were ordered by increasing value of the intensity of the tag. The ordered data were grouped into bins (subpopulations) of equal length *r* (number of cells) large enough to suppress intrinsic noise. After averaging the intensity data in each bin, the dimension of the resulting data array is reduced by a factor *r*. This coarsegrained process ensures reduction in intrinsic noise due to the Law of Large Numbers (cf. Supporting Information). Alternatively, noise filtering can be implemented by moving the average approach, preserving data dimensionality.

A similar strategy was employed for coarse-grained analysis of barcoded multi-drug experiments. Cell intensity data were clustered hierarchically based on the barcodes’ signals, and renormalized data were produced by averaging all optical parameters within each cluster (cf., Supporting Information).

Specifically, for siRNA experiments, after image segmentation, fluorescence of QD drug carriers and of GFP was assessed for individual cells. Each drug’s uptake was assumed to be proportional to the intensity of the corresponding QD carrier in each cell. The data were typically recorded as a 3-tuple {*QD*_1_, *QD*_2_, *GFP*}_*i*_ for each cell *i*.

We grouped cells into bins characterized by different QD combinations (in terms of fluorescence intensities {*QD*_1_, *QD*_2_}). For pairwise drug combination experiments, binning was achieved by partitioning the 2d space of the corresponding QD-tags’ fluorescence values into a finite mesh. The bins were designed to contain 30 or more cells each. We averaged a phenotypic marker (GFP intensity in our case) over each binned cell population.

### Cell Culture and Microscopy Imaging

The great majority of the experiments were conducted with the HeLa cell line. We used human pulmonary artery endothelial cells (HPAEC) for quantification of auranofin delivery. All experiments utilizing the point injection method were performed in confluent cell culture. The cells were grown on culture plates (microscopy slides) as follows: we used rectangular 4 and 8 chamber slides (Ibidi μ-Slides) for siRNA experiments, and either rectangular or circular slides for dye labeling experiments.

In experiments requiring single-cell imaging, cells were either ‘shrunken’ or completely detached using Accutase treatment to facilitate image segmentation. Homogeneous dye and QD labeling were performed according to the manufacturer’s (Thermo-Fisher) suggested protocol(s), typically at room temperature.

For point injection experiments, we used 10x dye concentration (compared to the homogeneous staining condition). Delivery was carried out by injection of ~ 0.05 – 0.1 cell culture volume locally. Cell imaging was performed with an inverted fully motorized Zeiss LSM-800 confocal scanning microscope utilizing 405, 488, 561, and 640 nm diode lasers for excitation of dyes and markers. Most of the microscopy experiments were performed using the 5x objective to capture larger optical fields necessary for statistical analysis.

Imaging data from single-cell experiments were processed as follows: cell segmentation was performed using bright-field and/or dedicated cytosolic markers. We employed Cellpose [18] for accurate cellular segmentation. For the auranofin co-delivery experiments, the ratiometric signal of the HyPer7 sensor was calculated as the ratio of cellular fluorescence intensities of the GFP channel excited by 488 and 405 nm laser sources.

Excitation and emission settings were chosen according to the dye manufacturer’s sug-gestions. In all QD experiments, we used a 405 nm laser for excitation with corresponding emission spectra filters. We tested fluorescent labels/markers individually to detect possible cross-channel leakage/overlap. This analysis was especially critical in dye co-delivery experiments where the degree of correlation may be strongly affected by channel leakage. To ensure channel “isolation,” we utilized dyes/fluorophores with well separated emission spectra for these experiments.

HeLa cells were maintained in Dulbecco’s modified Eagle’s medium (DMEM) containing 0.11 g/liter sodium pyruvate, 2 mM L-glutamine, 4.5 g/liter glucose, and 10% fetal bovine serum (FBS). Human pulmonary artery endothelial cells (HPAEC) were cultured at 37°C in modified EGM2 medium (Clonetics). Cells were maintained in continuous culture in an air-5% CO_2_ atmosphere at constant humidity and used within three to five passages. For fixation experiments, after aspiration of the culture media, cells were incubated with 10% formalin for 5 minutes at room temperature followed by a PBS wash. For cell suspension experiments, we detached HeLa cells with Accutase solution (Sigma-Aldrich), washed the cells with culture media, and kept them at either room temperature or 4°C in dPBS in non-treated plastic cuvettes for a short duration of dye(s) labeling (typically 1/2 hour).

In siRNA experiments, point-injection of two different QD nanoparticles carrying different siRNAs was performed at two different corners of the cell culture plate (chamber slide). Samples were incubated at 37° for 30 minutes. [Of note, no special precautions were taken to suppress the convection process, which facilitated the creation of concentration gradients.]

### siRNA design and delivery method

We used short-lived green fluorescent protein (d2EGFP) to ensure correlation between fluorescent GFP signal and mRNA target concentration in cells. The d2EGFP plasmid [19] was a gift from Derrick Rossi, Addgene #26821. We used siRNAs that were designed and experimentally validated by Dharmacon (cf., Accell eGFP Control siRNA) and the Sharp laboratory with binding strand sequences GCCACAACGTCTATATCAT (siRNA_1_) and GCACCATCTTCTTCAAGGA (siRNA_2_), respectively. The binding sites of siRNA_1_ and siRNA_2_ are separated by distance *d =* 153 nucleotides (in primary sequence). The msiRNA_1_ construct has a single nucleotide mismatch in the binding strand, GCCACAACG**G**CTATATCAT, and the separation distances to siRNA_1_ and siRNA_2_ were *d* = 0 and *d* = 153 nucleotides, respectively.

We utilized a fluorescent tag for each siRNA that was retained by the cell as follows. All siRNAs were designed with a cleavable disulfide linker (CL) attached to their 3’ end (Figure 5). We used streptavidin-conjugated Quantum Dots (QDs) as fluorescent tags for each class of siRNAs, creating uniquely labeled siRNA-QD pairs. Quantum Dots (QDs) are semiconductor nanoparticles, typically excitable by UV light, with fluorescence emission spectra that are fairly narrow. The emission peaks of QDs depend on the nanoparticle size, a property that makes QDs very useful for high-throughput microscopy measurements [20]. To facilitate siRNA-QD cellular delivery, we further functionalized the QDs with a cell penetrating peptide (CPP) [a nanopeptide (Arg)_9_] that can facilitate cell uptake through endocytosis or non-endocytotic mechanisms [21] (cf., Supporting Information for technical details).

### Coarse-Grained Image Analysis

In addition to course graining of single-cell data, we develop a similar approach with respect to raw images. We performed effective local averaging over multiple confluent cells by partitioning the entire image into tiles. The size of the tile (typically, ~ 50 x 50 microns) was estimated to contain at least a few cells on average.

Each tile in the image was replaced with an effective “pixel” characterized by the average fluorescence intensity of real pixels within the tile. The only segmentation we performed in this case was identification of large lacunae (cell-free regions) in the optical field, with exclusion of these regions from image analysis.

## Acknowledgement

The authors wish to thank Arvind K. Pandey for assistance with the HyPer7 probe, William M. Oldham for numerous insightful suggestions on experimental design, and Stephanie C. Tribuna for assistance in preparation of this manuscript. This work was supported, in part, by NIH grants HG007690, HL108630, HL155107, HL155096, and HL119145, and by American Heart Association grants D700382 and CV-19, and AHA957729.

